# Breakdown in seasonal dynamics of subtropical ant communities with land-cover change

**DOI:** 10.1101/2023.01.17.523860

**Authors:** Jamie M. Kass, Masashi Yoshimura, Masako Ogasawara, Mayuko Suwabe, Francisco Hita Garcia, Georg Fischer, Kenneth L. Dudley, Ian Donohue, Evan P. Economo

## Abstract

Concerns about widespread human-induced declines in insect populations are mounting, yet little is known about how land-use change modifies the dynamics of insect communities, particularly in understudied biomes. Here, we examine how the seasonal patterns of ant activity—key drivers of terrestrial ecosystem functioning—vary with human-induced land cover change on a subtropical island landscape. Using trap captures sampled biweekly from a biodiversity monitoring network covering Okinawa Island, Japan, we processed 1.2 million individuals and reconstructed activity patterns within and across habitat types. Forest communities exhibited greater variability than those in more developed areas. Using time-series decomposition to deconstruct this pattern, we found that ant communities at sites with greater human development exhibited diminished seasonality, reduced synchrony, and higher stochasticity compared to those at sites with greater forest cover. Our results cannot be explained by variation in either regional or *in situ* temperature patterns, or by differences in species richness or composition among sites. We conclude that the breakdown of natural seasonal patterns of functionally key insect communities may comprise an important and underappreciated consequence of global environmental change that must be better understood across Earth’s biomes.

## Introduction

Insects comprise 95% of described terrestrial animal species on Earth and are key drivers of a multitude of ecosystem functions and services, including pollination, food provisioning, pest control, water filtration, carbon sequestration, and decomposition (Schowalter, 2013). Mounting evidence about global insect declines due to human-induced environmental change presents a palpable threat to a sustainable future for humanity (Goulson, 2019; Wagner, 2020). However, our understanding of global insect declines is in general based on year-to-year trends that overlook intra-annual change in seasonal patterns, which can only be detected by sampling at fine temporal resolutions (Tonkin et al., 2017). For example, the majority of studies on community temporal variability compare measurements between years and rarely within them (e.g., Cottingham et al., 2001; de Mazancourt et al., 2013; Olivier et al., 2020; Tilman et al., 2006), meaning that modification of seasonality due to anthropogenic factors may be overlooked. When community seasonal patterns are altered by human pressures, ecosystem functions and services can be degraded (Ross et al., 2021; Stevenson et al., 2015). For this reason, a better understanding is needed of how fine-scale temporal patterns of insect communities are affected in this era of accelerating global change.

Land-cover modification is known to have negative effects on insect diversity (e.g., Corro et al., 2019; Oliver et al., 2016; Senapathi et al., 2015). Corresponding effects on temporal community dynamics such as seasonality are, however, less well-documented. There is some evidence that human development can erode seasonality of aquatic insects (Castro et al., 2018), bees (Hung et al., 2017), bugs and leafhoppers (Knop, 2016), and butterflies (Uchida et al., 2018). For most of these studies, loss of seasonality is linked to biotic homogenization, whereby communities become more similar and generalist due to replacement of native species with exotics (McKinney, 2006). As maintenance of natural seasonal patterns is associated with more reliable provision of ecosystem functions and services (Stevenson et al., 2015), it is important to know whether native species that persist in communities affected by human development retain their seasonality. This question still remains unanswered, especially for understudied biomes such as the tropics and subtropics that are undergoing particularly rapid land-cover change (Janzen & Hallwachs, 2021), and also for islands, which have restricted land area, frequent introductions of alien species, and exposure to increasingly extreme weather events (Russell & Kueffer, 2019).

Here, we examine the effects of anthropogenic land-cover change on the fine-scale temporal variability of ant community activity. Ants are a dominant insect group in the many environments they inhabit around the world, underpinning a broad spectrum of ecological interactions and playing a major role in regulating ecosystem function (Hölldobler & Wilson, 1990). This makes them ideal bioindicators for ecosystem change (Andersen & Majer, 2004). However, very little is understood about how land-cover change affects ant community seasonality in less seasonal biomes outside the temperate zone, though it is known that ants do exhibit significant seasonality in tropical and subtropical regions (Basu, 1997; Samways, 1990).

To address this, we used high-resolution (biweekly) monitoring data on ant activity in Okinawa Island in southern Japan to examine whether, and how, the temporal variability of ant communities varies with anthropogenic land-cover change on a heavily populated subtropical island. We then determined the temporal components of variability (i.e., seasonality, trend, or stochasticity) that contribute the most to differences across the land-cover gradient (Fig. S1) and examined the importance of other factors potentially influencing patterns in community variability. Finally, we examined the responses of native and non-native species separately to determine their respective contributions to community dynamics.

## Materials and Methods

### Study sites and land-cover data

Our study focuses on the subtropical main island of Okinawa in the southern Ryukyu archipelago of Japan. The island has a historical land-cover gradient, spanning from the minimally developed north to the more urban south (Fig. 1). Okinawa’s extensive Yambaru Forest in the north was recently recognized as part of Japan’s newest Natural World Heritage Site for its high endemism and biodiversity. Although the island is only 0.3% of Japan’s total land area, it provides habitat to more than one-third of the country’s ant species (Terayama et al. 2009; Terayama et al. 2014). The few studies of ant communities in Okinawa include biodiversity surveys (e.g., Ito et al., 1998) and comparisons between native and alien ant populations in Yambaru (Katayama & Tsuji, 2010; Suwabe et al., 2009; Yamauchi & Ogata, 1995). This study, the first to sample island-wide ant diversity and temporal patterns in Okinawa, uses data from a biodiversity monitoring system covering the land-cover gradient of the island, administered by the Okinawa Environmental Observation Network (OKEON) Churamori Project (https://okeon.unit.oist.jp/).

**Figure 1.**
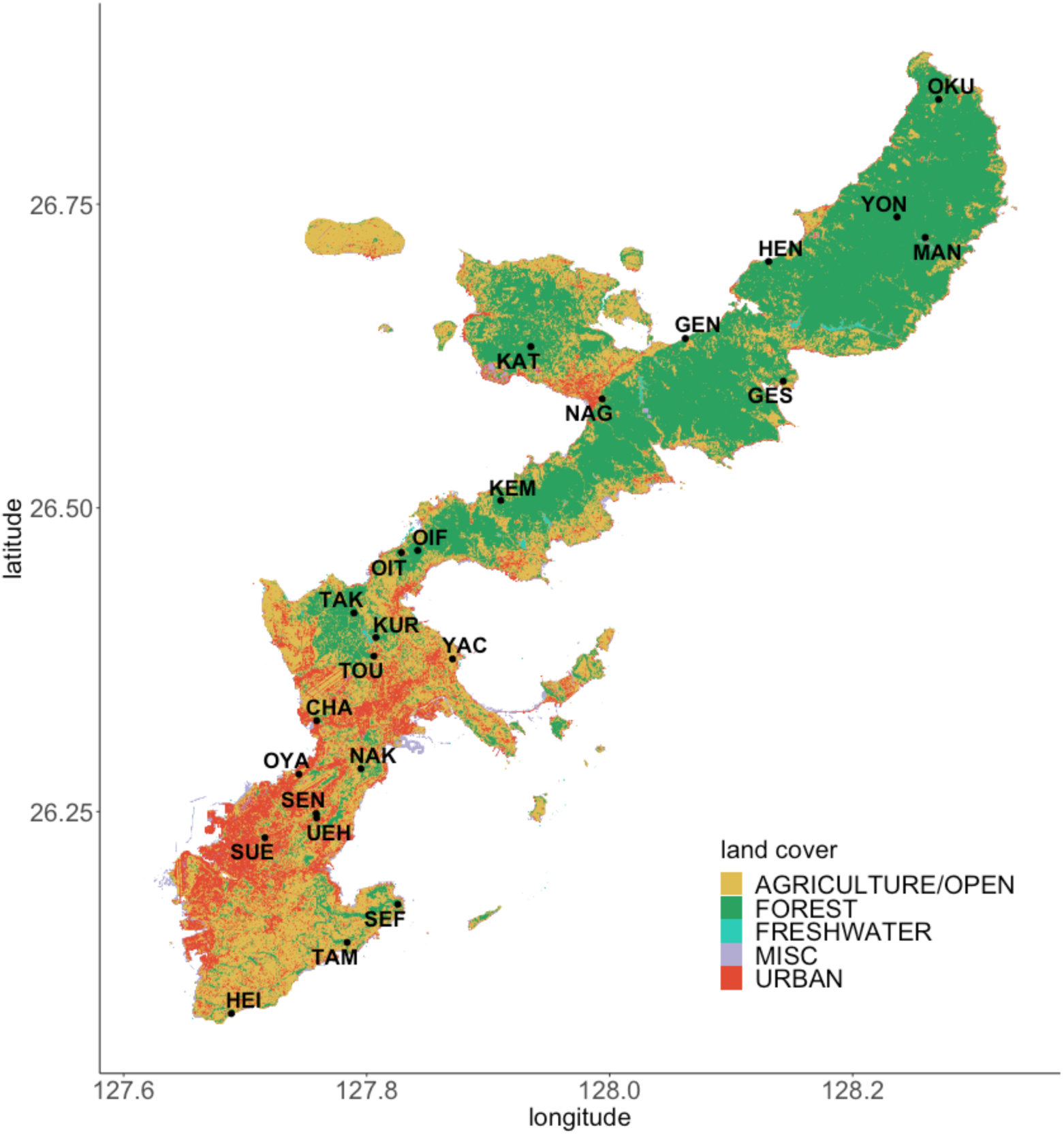
Major land-cover classes of Okinawa Island with locations of the OKEON monitoring stations, which cover the full gradient of human development across the island. Ants were sampled from three stations at each sampling site biweekly from 2016– 2018 (total of 52 sampling periods).

We used a land-cover map for Okinawa (year 2015) to calculate representative land-cover values for the 24 OKEON monitoring sites. The sites (pairwise distances: minimum = 389 m, mean = 33 km, maximum = 102 km) are located in or near forested areas across multiple land-cover types varying from broadly contiguous forest to highly agricultural or urban areas (Fig. 1). To characterize each site, we calculated the proportion of seven main land-cover classes (forest, agriculture, urban, grass, sand, freshwater, miscellaneous) around circular buffers with radius 1 km (see Ross et al., 2018 for details). Buffers with this size were chosen to characterize representative land-cover values for areas surrounding sites without including regions that may have resident unsampled colonies. Sites near the coast included ocean area within the buffer, so we used relative proportions for all sites that considered only land area. As correlations were high between the main land-cover classes of interest (that is, forest, agriculture, and urban; Table S1), we used principal component analysis (R function *prcomp*) after applying an arcsine transformation to the proportions to improve normality (e.g., Van Buskirk, 2005) and retained the first and second axes (variance explained: PC1 = 81% and PC2 = 11%) for use as explanatory variables. PC1 represents the forested (high) to developed (low; urban and/or agriculture) gradient, while PC2 represents the rural (high; agriculture and/or grass) to urban (low) gradient (Fig. S2). All analyses were conducted in R (R Core Group, 2021), and spatial analyses were conducted with R packages *sf* (Pebesma, 2018) for vector data and *raster* (Hijmans, 2021) for gridded data.

### Ant activity data

We sampled worker ant activity biweekly with Sea, Land, and Air Malaise (SLAM) traps for two years and identified samples to species level, resulting in 1,378,324 classified individuals.

Although Malaise-type traps are often used to catch flying insects, they also can be used to catch a variety of non-flying insects (Uhler et al., 2022), including a wide diversity of worker ants (Economo & Sarnat, 2012; Longino & Colwell, 1997). We chose to use SLAM traps as ground pitfall traps tend not to work in moist forest environments like Okinawa, as they quickly fill with snails, and Winkler extractions are too destructive to employ on a biweekly repeated basis. Like any entomological survey method, SLAM traps provide a biased sample of the community, in this case towards walking insects that climb on low vegetation. However, this is a consistent bias and no different from any trapping or detection method in ecology that targets a stratum of an ecosystem (e.g., leaf litter, canopy, soil, etc.). Our goal here is to examine differences in community dynamics across habitats, not estimate absolute abundances of species in the ecosystem. We sampled one SLAM trap (Large BT1005, MegaView Science Co., Ltd.) at each of three stations per OKEON site (pairwise distances within sites: mean = 90 m, minimum = 19 m, maximum = 195 m) every two weeks from March 2016 to March 2018 (further details in Supporting Information A). The stations are predominantly deployed under forest canopy, though several are in nearby open areas. This sampling method mainly targets terrestrial browsing species, though does occasionally catch arboreal and subterranean individuals. Insect count data sampled with passive traps are usually interpreted as a measure of activity, or the rate at which individuals intersect a point in space, rather than direct estimates of abundance, though differences in abundance are generally reflected in ant activity data (Kaspari et al., 2022a).

We additionally categorized all species by invasive status as either native, alien, or uncertain based on expert opinion and biogeographical databases (Guénard et al. 2017, Janicki et al. 2016, Terayama et al. 2009, https://www.antwiki.org/, https://www.antweb.org/). Native refers to species with enough data to establish historical occupancy in Okinawa without human-aided dispersal from other regions. Alien refers to species that definitively have native ranges in different regions and have dispersed to Okinawa via human activity. Uncertain species lack sufficient data to make such determinations.

Some prevalent species had extreme count outliers, such as *Technomyrmex brunneus* (representing 2-week counts greater than 5,000), likely caused by unusual events such as moving to nests nearby or intense foraging at specific food sources rather than overall trends in activity patterns. As these rare events could have a large effect on variability estimates, we limited count values to a conservative threshold of 500 individuals for a given species at a station within a two-week period. We also examined changes in our results for a range of thresholds to check for sensitivity of results. For each species during each sampling period, we summed counts over the three stations per site to determine site-level activity (Fig. 2), and over all sites per land-cover group for grouped species-level activity. We then calculated total site density as the sum of all counts per site.

**Figure 2.**
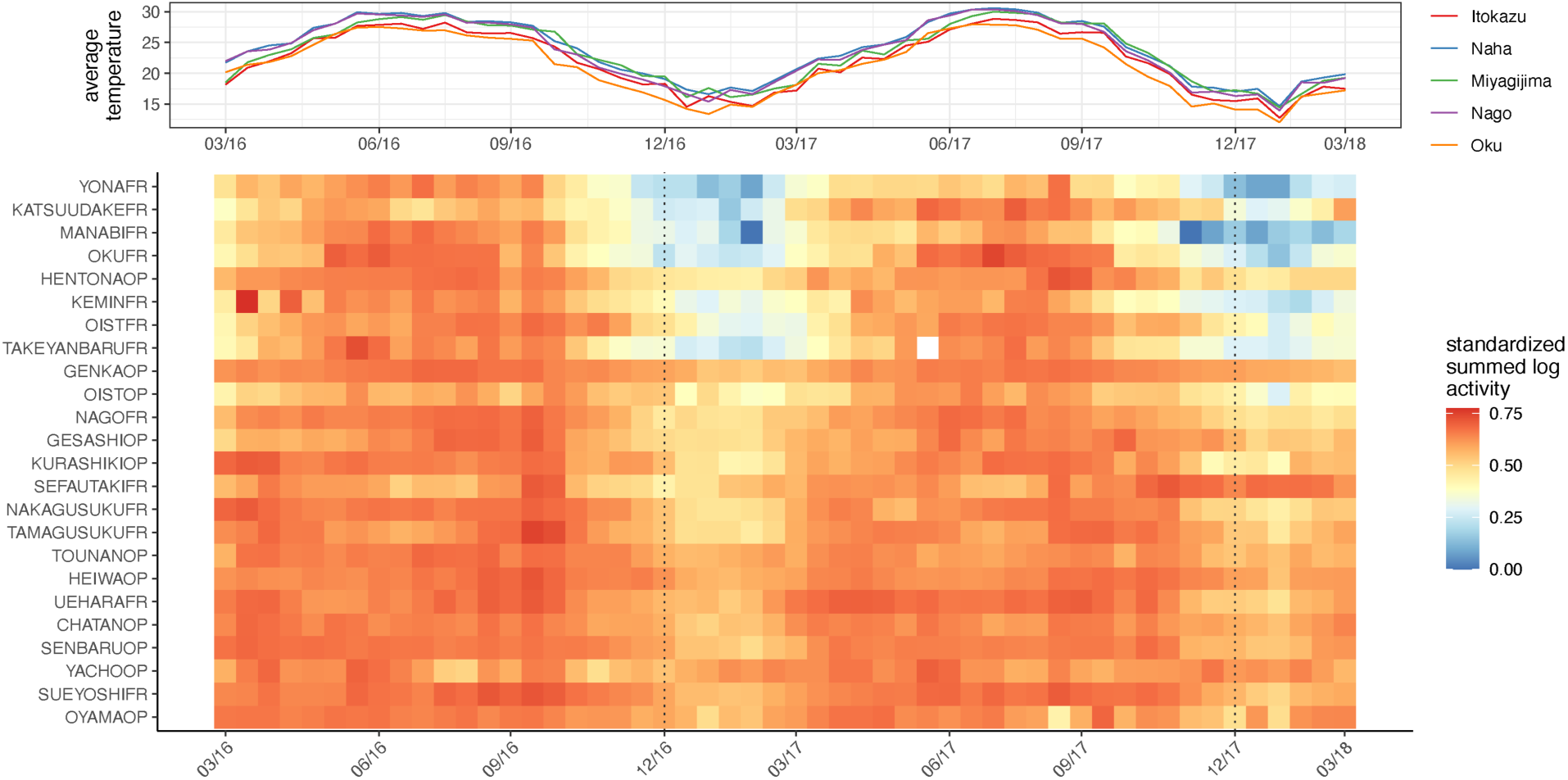
Heat map showing the standardized ant activity (i.e., logs of summed site counts divided by the log of the grand total count) over two years in Okinawa. Mean temperature measurements from the Japan Meteorological Agency across the island, ordered from south to north, are included for reference. Heat map sites are ordered by values of land-cover PC1, explaining the anthropogenic stress gradient (top: more forested, bottom: more urban). Dotted lines show the last sampling period before the new year.

Sampled abundance and variability can be correlated due to sampling error alone because of mean-variance scaling (McArdle & Gaston, 1995). For example, even if species dynamics are identical across two sites, if we sample fewer individuals from site A than site B (e.g., due to effects of sampling station placement), site B may have a higher temporal variability simply because its counts are higher. While in some sense this could reflect the actual variability of species encounters at specific locations (that is, at the stations), we wanted to disentangle differences in community variability from those attributable to sampling error. Thus, we rarefied the data from all sites so that the total count (sum of all samples across the two years) for each site equals the total count at the site with the least abundance (Fig. S1, Supporting Information B; Gaston & McArdle, 1994). Finally, to determine whether species richness differences could drive any observed differences in temporal variability, we calculated site-specific total species richness, and the richness of each invasive status group from the observed and rarefied data, making extrapolations to account for incomplete sampling using Hill numbers (*q* = 0) with the R package *iNEXT* (Hsieh et al., 2016).

### Temporal variability and time-series decomposition

We calculated two aspects of temporal variability: functional and compositional (Supporting Information C). Functional variability refers to activity changes at the aggregate scale (e.g., Hillebrand & Kunze, 2020) and is calculated as the coefficient of variability (ratio of the standard deviation to the mean) of total ant counts across all species per site (McArdle & Gaston, 1995). Compositional variability estimates the variation in community composition over time and is calculated as temporal beta diversity (Legendre & De Cáceres, 2013) using the R package *adespatial* (Dray et al., 2022), which additionally records species’ individual contributions to beta diversity. As we use ant count data, compositional changes would thus represent changes in relative activity rates of different species in the local species pool, which in turn is a function of the abundance and patterns of behavior (Kaspari et al., 2022a).

Next, we used temporal decomposition models to determine the relationships of temporal variability components with observed patterns. Time series can be decomposed into components describing different underlying temporal patterns in the data: “seasonal” processes that repeat cyclically over some temporal scale (i.e., phenology), “trend” that describes directional change, and “remainder” that represents any residual variation not captured by the other components (i.e., short-term stochastic fluctuations; Hyndman & Athanasopoulos, 2021). A community with high temporal variability may have strong cyclical patterns, increasing or decreasing trends, high stochasticity, or some combination of these. For temporal datasets spanning many annual cycles, models that can estimate complex seasonal responses (e.g., wavelet analysis; Tonkin et al., 2017) are often employed, but for studies spanning few annual cycles at a fine temporal resolution, simpler models with fewer assumptions and problems with overfitting are preferable. We estimated additive temporal components for the ant activity data by fitting time-series linear models with temporal predictors using the R package *fable* (O’Hara-Wild et al., 2021). As our time-series was short, we used simple predictors consisting of a linear trend and a seasonal signal approximated with an annual Fourier term with the simplest maximum order (*K*) of 1—this models seasonality as a sine wave (Hyndman & Athanasopoulos, 2021). We fitted this model to the rarefied data at both the site-level and species-level for land-cover groups, then decomposed the count value at each time-step. For each site, we measured the absolute variance of each temporal component, but also the relative component variance, defined here as the individual component variance divided by summed variance of all components (Supporting Information D).

Finally, as spatial autocorrelation between sites did not affect results (Table S3; Supporting Information E), we used simple linear models to estimate relationships between land cover and our focal metrics of temporal variability. Specifically, we fitted models for total ant counts (before rarefaction), functional and compositional temporal variability, and absolute and relative variances of time-series components. To explore differences between native and alien species, we also fitted models separately by invasive status for site richness and summed species’ contributions to beta diversity. We performed model selection on linear combinations of the predictor variables land-cover PC1 and PC2 using the Akaike Information Criterion corrected for small sample sizes (AICc).

### Assessing effects of differences in temperature and community composition on observed seasonality patterns

To determine if other differences among sites may be responsible for any observed variation in ant community activity and relationships with land cover, we additionally examined differences in 1) regional and site-level temperature and 2) tested whether seasonality differences persisted after standardizing community composition between land-cover groups. For (1), we downloaded regional climate data for the collection period from the Japan Meteorological Agency (JMA) database (http://www.jma.go.jp/en/amedas_h/map65.html, accessed 04/01/2020) and calculated temperature means and extremes from the six climate stations that collect temperature data (Miyagijima, Itokazu, Naha, Ashimine, Nago, Oku). We also collected *in situ* site-level data throughout the collection period at one station per site to characterize local air and soil temperature patterns (WatchDog 2900ET Weather Station 1.5 m above ground and SMEC WaterScout 300 Soil Sensor 10 cm below ground, Spectrum Technologies). With the same workflow as for the ant activity data, we calculated absolute variance of seasonality for regional and site-level temperature time-series, then estimated relationships with land-cover PC1 using linear models.

For (2), we conducted analyses to test whether differences in community composition alone, without variation in seasonality within species, could be driving the observed patterns in seasonality, or alternatively if habitat-dependent dynamics of species occurring in both habitat types were responsible. We used land-cover PC1 to partition the sites into two groups with eight sites each—forested and developed (comprising sites with more agricultural and/or urban land cover)—and retained only those species shared between groups and with total counts greater than 100. We excluded the eight sites with intermediate values of PC1 to better represent the most characteristic sites for each land-cover group. To make balanced comparisons between land-cover groups, we rarefied each species’ activity data to the minimum count of that species between groups (Fig. S1, Supporting Information B). To determine whether community composition affected our results, we fitted time-series linear models to each species in the land-cover groups and calculated the relative variance of their seasonality (forested versus developed sites). As differences between the land-cover groups were not normally distributed (Shapiro-Wilk test; *p* = 0.011), we used the nonparametric Wilcoxon signed-rank test to determine whether the same species between groups exhibit significant differences in relative seasonality variance. Additionally, to determine how temporal mismatch changes with invasive status, we calculated the synchrony of seasonality for each invasive status between land-cover groups and standardized them to the same scale. We used the R package *codyn* (Hallett et al., 2016) to calculate synchrony using the method described by Loreau & De Mazancourt (2008). We then compared seasonal curves to the average seasonality curve for each land-cover group to visualize the degree of temporal mismatch by invasive status.

## Results

Our sampling for 2016–2018 resulted in the recovery of >1.2 million individuals (before thresholding) and 16,595 unique count records at the site-level across 91 ant species. Total count was less than 5 individuals for 21 species and greater than 10,000 individuals for 15 species, with a minimum of 1 (*Aenictus ceylonicus*, *Crematogaster* c.f. *matsumurai*, *Ectomomyrmex sp.*, *Erromyrma latinodis*, *Hypoponera* sp., *Leptogenys confucii*, *Strumigenys hirashimai*, *Strumigenys mazu*) and a maximum of 259,180 (*Tetramorium bicarinatum*). The pattern of ant activity dropping in the winter and rising in the summer qualitatively diminished as human disturbance increased (Fig. 2). Total ant activity (after thresholding to 500) had a nearly 23-fold difference between the site with the minimum count (Yona Forest, *n* = 3616) and that with maximum count (Sueyoshi Forest, *n* = 82,781). The natural log of total density had a strong negative relationship with land-cover PC1 (R^2^ = 0.71, *p* < 0.001), indicating that ant activity overall was higher in developed areas than forested areas (Fig. 3). Rarefaction resampled all sites to the minimum site count before analysis.

**Figure 3.**
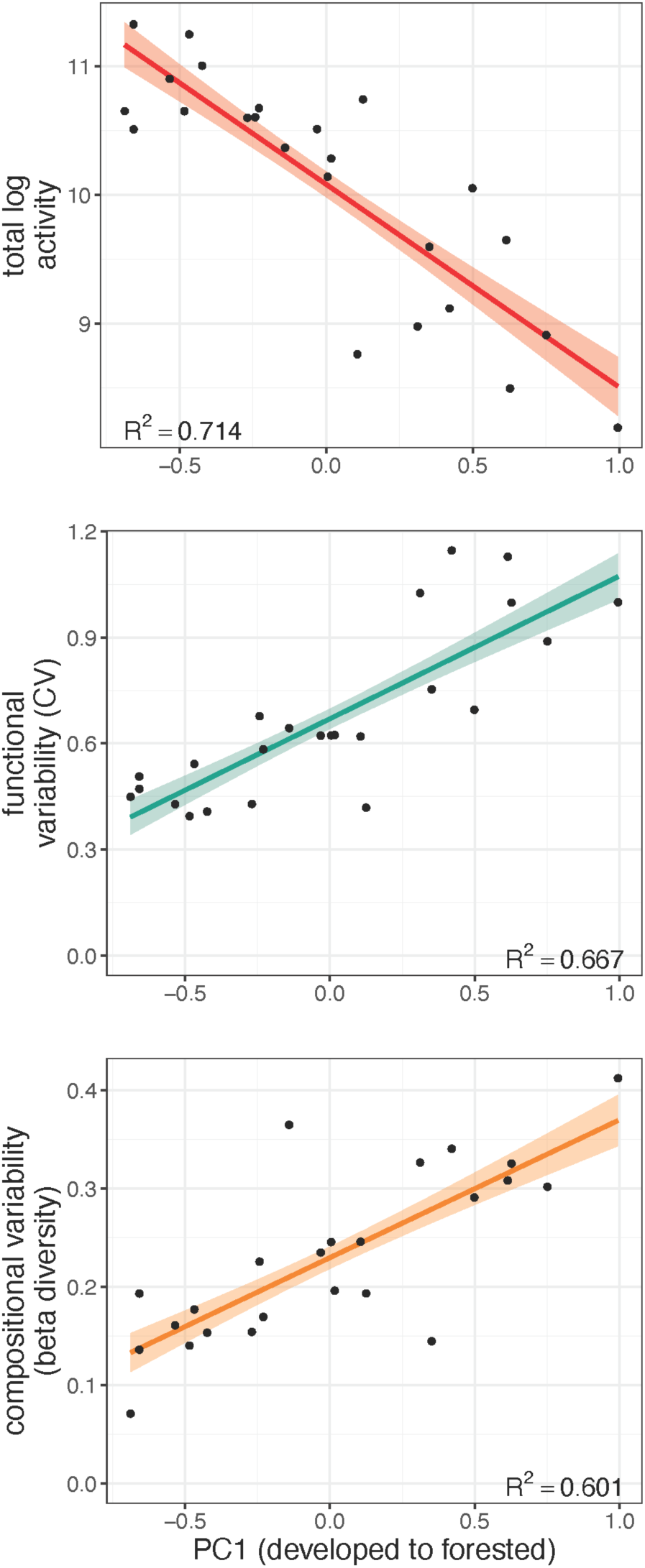
Trends in total log activity, functional variability, and compositional variability for ants across a gradient of anthropogenic impact. Relationships with PC1, explaining the anthropogenic stress gradient (low: more developed, high: more forested). Total log ant activity (before rarefaction) correlates negatively with PC1 while indices of temporal variability (functional: coefficient of variation, compositional: beta diversity) correlate positively.

The linear models suggested strong relationships with the land-cover PC1 for most metrics of temporal variability (Figs. 3, 4). Based on model selection via AICc, all models included PC1 as the sole predictor except the relative variances of seasonality and stochasticity, which also used PC2 (Table S3). Functional and compositional temporal variability both had strong positive relationships with PC1 (functional: R^2^ = 0.67, *p* < 0.001; compositional: R^2^ = 0.60, *p* < 0.001; Fig. 3), and these two variabilities also had a strong relationship to each other (R^2^ = 0.62, *p* < 0.001; Fig. S3). The species’ contributions to beta diversity (Fig. S4) that explained individual species’ impacts on compositional variability were also positively correlated with PC1 for native species (R^2^ = 0.40, *p* < 0.001), but uncertain (R^2^ = 0.28, *p* < 0.01) species had a weaker, negative relationship (alien species showed no relationship). Absolute seasonality and stochasticity variance had positive relationships with PC1, though that for seasonality (R^2^ = 0.57, *p* < 0.001) was stronger than that for stochasticity (R^2^ = 0.15, *p* < 0.05; Fig. 4). On the other hand, relative seasonality variance (R^2^ = 0.23, *p* < 0.05) was positively related to PC1, while relative stochasticity variance (R^2^ = 0.36, *p* < 0.01) had a negative relationship (both had *p* > 0.05 for PC2; Fig. 4). All linear regression results are found in Table S3.

**Figure 4.**
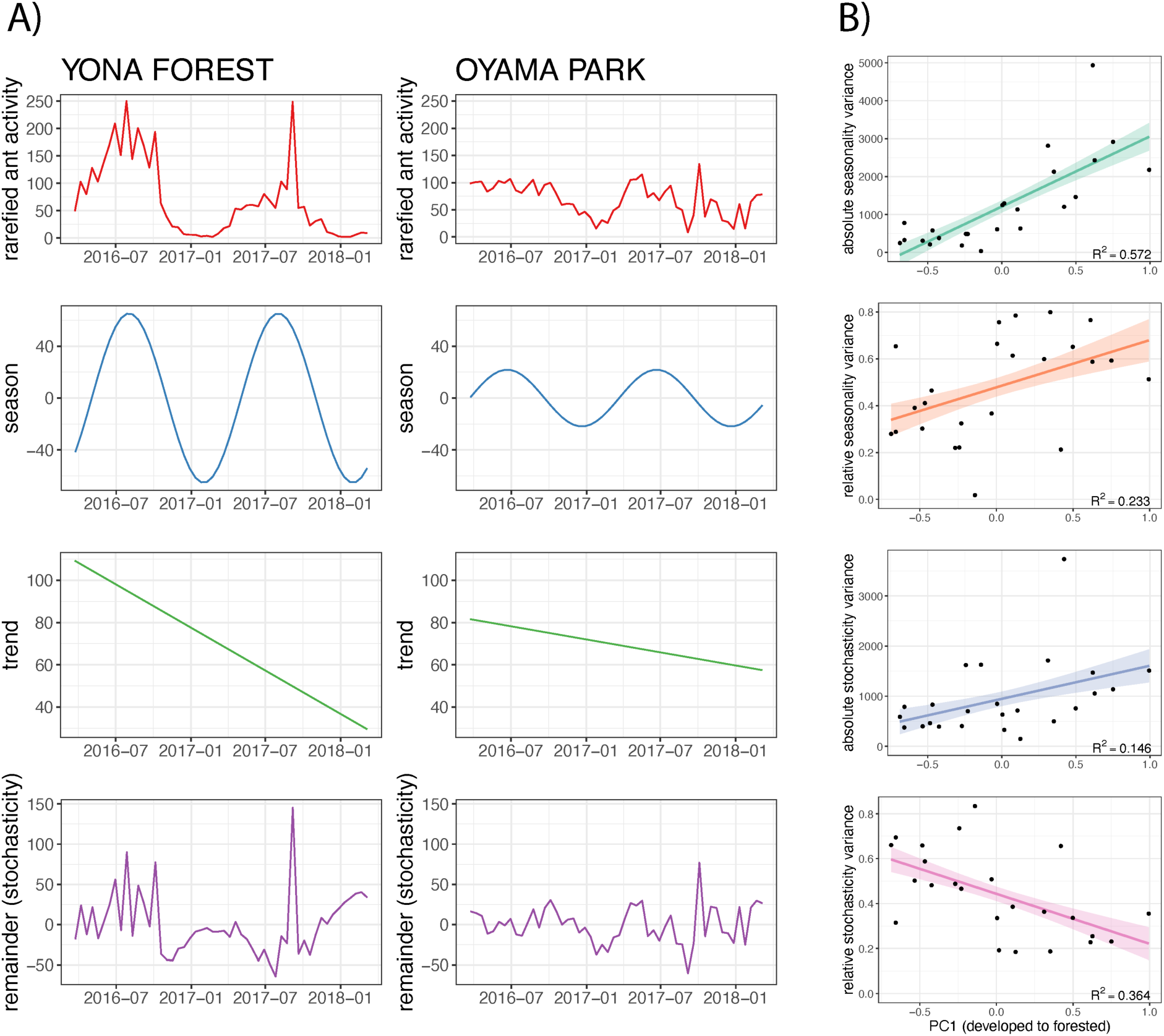
A) Time-series decomposition of rarefied ant activity for the sites with the most forest (Yona Forest) and most human development (Oyama Park) within a 1 km buffer. B) Relationships between absolute and relative variance of seasonality and stochastic temporal components and PC1 across all 24 sites, explaining the anthropogenic stress gradient (low: more developed, high: more forested). Relative variance here is calculated as individual component variance divided by summed variance of all components.

We found no evidence that differences in species richness, temperature at different scales, or species composition are candidates as drivers of observed differences in ant community temporal variability. We found no relationship between PC1 and total species richness (either observed or extrapolated), and a weak correlation with rarefied richness (R^2^ = 0.15, *p* < 0.05) due to the loss of rare species during the rarefaction process, though we consider this unlikely to bias our results (Fig. S5). We did find strong positive relationships with PC1 for native richness values (observed: R^2^ = 0.47, *p* < 0.001; extrapolated: R^2^ = 0.35, *p* < 0.005; rarefied: R^2^ = 0.61, *p* < 0.001) indicating that more native species can be found in the more forested sites, and weaker negative relationships with richness for uncertain (observed: R^2^ = 0.32, *p* < 0.005; extrapolated: R^2^ = 0.33, *p* < 0.05; rarefied: R^2^ = 0.33, *p* < 0.05) and alien species (observed: R^2^ = 0.29, *p* < 0.005; extrapolated: R^2^ = 0.20, *p* < 0.05; rarefied: R^2^ = 0.20, *p* < 0.05) (Table S3, Fig. S5).

Moreover, we found no clear patterns in the absolute variance of regional temperature seasonality from JMA climate stations spanning the island from north to south, nor did we find relationships between the absolute seasonality variance of *in situ* air or soil temperature and PC1 (Fig. S6). Only two sites had exposed sensors that were not located below canopy cover (Oyama Park and OIST Open [OYA and OIT, respectively, in Fig. S6]), and these had higher absolute variances than the other sites. Concerning the tests of different threshold sizes, although we observed some differences, our results remained similar (Supporting Information F, Fig. S7).

Lastly, individual species showed differing seasonality between land-cover groups (more forested and more developed). Comparisons revealed that, in general, the same species have a higher relative seasonality variance at forested sites than at sites with more human development (*p* < 0.005), though the alien species *Tetramorium lanuginosum* was a notable exception (*Tlanu* in Fig. 5). While seasonality was stronger in the forested group for all species, when separated by invasive status we observed higher synchrony in the forested group for native (*n* = 11, forested: 0.75, developed: 0.58) and uncertain species (*n* = 8, forested: 0.5, developed: 0.28), but the opposite pattern for alien species (*n* = 4, forested: 0.73, developed: 0.86) (Fig. 5).

**Figure 5.**
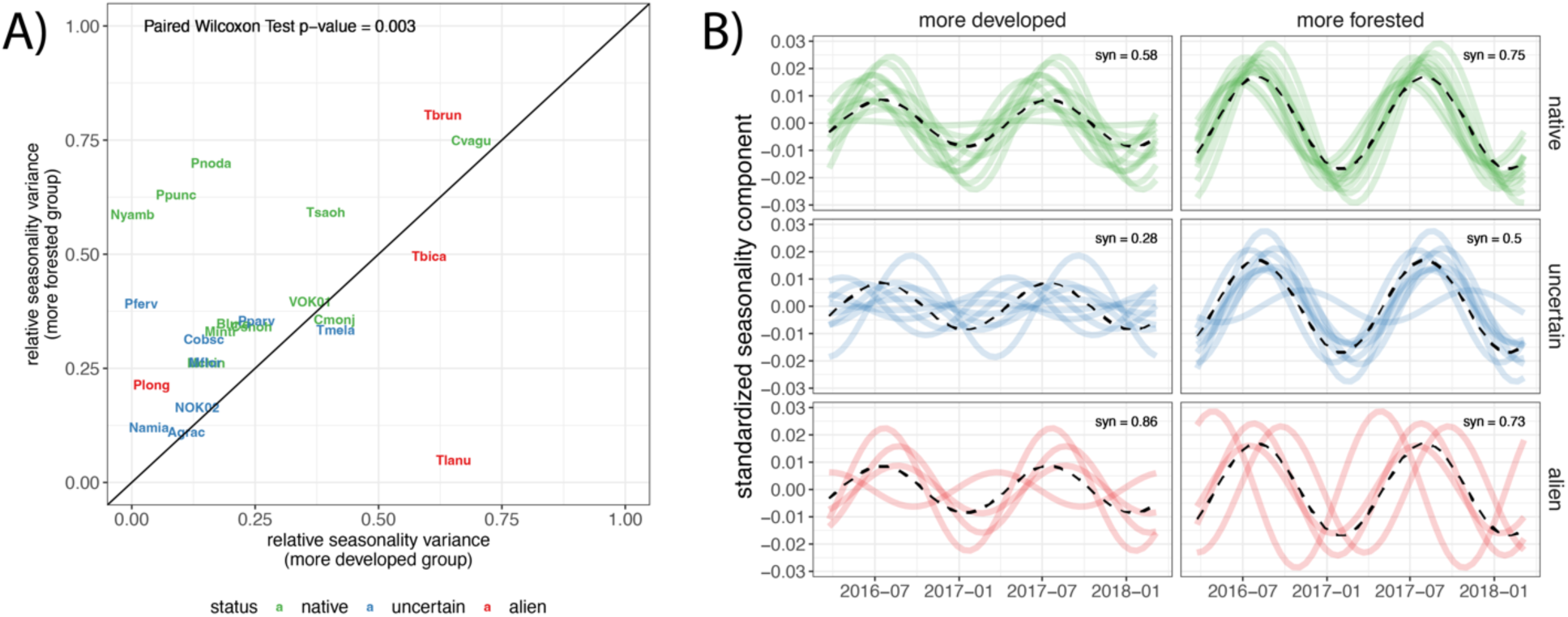
A) Relative seasonality variance calculated per ant species in two representative land-cover groups each containing eight sites: more forested and more developed. Only species shared between groups (abbreviations of those found in Table S2) and with total counts ≥100 were retained, whereupon species counts were rarefied to the minimum between groups. On average, species have a higher relative seasonality variance in the forested group (paired Wilcoxon Test; p = 0.003), and no apparent patterns exist regarding alien status. B) Seasonal component time-series for each species (lines) found in the more forested and more developed groups, standardized by total species activity and separated by invasive species status. The dashed black line shows the mean standardized seasonality across all species per land-cover group, showing much greater temporal mismatch in ant communities at sites with greater human development.

## Discussion

Our study sheds light on how ant community activity dynamics vary across a gradient of human-induced land-cover change in a subtropical island environment. Notably, we show that land-use changes the temporal activity dynamics of the ant community by degrading the seasonal cycle.

Our results complement an extensive number of recent studies focusing on the temperate zone that examine change of insect populations over time (Hallmann et al., 2017; Hung et al., 2017; Kaspari et al., 2022a; Seibold et al., 2019; Uhler et al., 2021), including those few with enough intra-annual sampling to examine changes to seasonality (Hung et al., 2017; Kaspari et al., 2022b). However, those studies from the temperate zone are from ecosystems with dramatic seasonal variation in temperature that essentially forces seasonality in activity patterns. These are not representative of huge fractions of the subtropics and tropics where temperature varies meaningfully, but does not enforce a shutdown of the insect community. Thus, the effect of environmental degradation on seasonality of insect populations in such environments is still not well understood (Kishimoto-Yamada & Itioka, 2015).

Our study fills this an important gap by showing that ants, an ecologically dominant and functionally relevant group, completely change their activity patterns across the ecological disturbance gradient. We found that temporal variability diminishes for the subtropical ant communities of Okinawa Island as land cover transitions from forested to more developed, and that this is related to a breakdown in seasonality. Forested sites were more variable overall, but their resident ant communities were more seasonal and synchronous, while community variability for developed sites (i.e., urban and agricultural) was characterized by stochastic variation. Although native ant richness was higher in forested sites, we determined the observed differences in seasonality across the land-cover gradient were not due to differences in regional or local temperature patterns, nor to species composition among sites. Compositional differences between communities in different land-cover types, such as the prevalence of one or more highly seasonal species, could drive observed relationships between community seasonality and land cover. However, we found that populations of the same species occupying forested sites were more seasonal than those with more human development. Further, when we separated species by invasive status, we found that the strong seasonal patterns we observed in the forest were more synchronous for native than alien species, though both synchrony values are reasonably high and the considerably smaller sample size for alien species should be taken into consideration.

We found a positive relationship between functional and compositional temporal variability, meaning that those sites with communities that varied more as a whole tended also to have higher total variation of the individual species in the local pool. The relationship between functional and compositional variability can range from positive to negative depending on the system and exposure to disturbance (Hillebrand et al., 2018; Ross et al., 2022; White et al., 2020). Two characteristics of more forested sites may help explain this positive relationship. The first is that more forested sites had higher variability in general at the biweekly timescale. The second is that, at more forested sites, native species had higher richness and the highest contributions to beta diversity. Sites with high richness that include substantial numbers of alien species can have community dynamics that differ considerably from similar sites in which native species dominate (Krushelnycky & Gillespie, 2008; Sanders et al., 2003), thus including alien species in richness estimates can affect how diversity-stability relationships are interpreted (e.g., Moore & Olden, 2017).

Our results raise the question of what mechanisms drive these patterns. We outline several hypotheses for why seasonal variation in activity could vary across the land-cover gradient. First, there is a possibility that ants are simply responding to the temperature they experience on a physiological level, reflecting temperature and microclimatic variation across habitats. Ants are well-known to be sensitive to temperature variation, with foraging limited to certain temperature ranges (Bernstein 1979, Stuble et al., 2013). Regional climatic variation across the island is very modest, and with no consistent differences in seasonality. Further, we found no relationships between the seasonal variance of *in situ* air and soil temperature and the main land-cover gradient PC1. However, differences in canopy openness, vegetation, and urbanization (via heat-island effects, etc.) may create conditions that are relevant for promoting and depressing ant foraging, and the prevalence of such conditions might be higher in more degraded habitats.

Comprehensive measurements with small sensors at various habitat strata for representative sites could help elucidate whether this phenomenon could differ across land-cover types. Another possibility is that ants are responding to resource availability patterns. In this scenario, it is possible that resources are more seasonal inside the forest rather than more human-impacted areas. This could occur, for example, because of anthropogenic food sources that are more static throughout the year, or because of differences in phenology of vegetation inside or outside the forest (Penick et al., 2015). A related question is whether these differences in seasonality are limited to ants or are observed across the arthropod community. Were arthropod seasonality responding more generally to land cover, this would provide a resource pattern that could affect ant activity. Related to this, it is unclear why there seems to be little seasonal niche differentiation for ant species in forested areas, which have activity patterns that follow temperature quite closely in synchrony. One possibility is that species become most active at different times during the same warm season to avoid competition—we saw some evidence of this in the standardized seasonal curves (Fig. 5). This could be due to the inability to achieve high activity levels during cooler months due to physiological limitations, which may be absent in more developed environments with anthropogenic temperature refugia. Other potential explanations relating to differences in the ecology or physical environments across the two habitats likely exist, but more targeted research is needed to address this question.

Our results come with a few caveats. First, the fact that our data represents only two years is a limitation of our study. However, our sampling was very dense within those two years (24 sites, 72 traps, 52 samples per trap, >1.2 million individuals observed), and the analytical methods we used were specifically geared toward decomposing short time series into periodic, trend, and stochastic components. Thus, we are confident that our conclusion that seasonality depends on land cover is robust for the two years involved. It seems unlikely that these two years would be outliers, with seasonality greater in forest habitats in some years but not others, however we cannot rule this out. In any case, the broader conclusion that activity dynamics depend on land cover would still be valid. Finally, it remains to be seen whether our results for ants foraging in low vegetation generalize to the entire insect community and across habitat strata.

We measured variability over two years, but there may be other types of variability that only manifest on longer timescales and are also correlated with land cover. For example, introduced ants are known to have boom-bust dynamics when they reach new localities (Lester & Gruber, 2016). Over longer timescales, communities in more human-dominated areas may prove to be more dynamic with successive shifts in the community. Indeed, we witnessed such an event in our two-year period at one of our sites (Sefautaki Forest), where *Pheidole megacephala*, a notoriously impactful invasive ant, arrived and quickly reached high abundance, raising variability of this site beyond what was typical of others with similar land-cover characteristics. Recent research in Yambaru Forest has documented resilience of native ant communities to disturbance based on surveys of invasions at developed areas after regrowth (Shimoji et al., 2022), so this site too may eventually recover to its previous structure. Thus, successive regime shifts in these more disturbed communities could occur over longer timescales, but only monitoring over more years would reveal such dynamics.

## Conclusions

This study provides novel insight into the seasonal dynamics of subtropical insect communities across differing levels of human development. Importantly, it is the first study to our knowledge to measure seasonality of ant activity with high-frequency sampling across a human disturbance gradient. The results of our study, which link anthropogenic disturbance in the form of agricultural and urban development to a reduction in intra-annual temporal variability due to weakening seasonality, support the growing evidence that human development reduces natural seasonal patterns (Hung et al., 2017; La Sorte et al., 2014; Leveau, 2018; Uchida et al., 2018).

Although this study focuses on correlative relationships and thus cannot fundamentally demonstrate causal mechanisms, it does provide compelling evidence that continuing habitat loss and fragmentation can lead to increasing homogenization of insect communities in parts of the globe with the highest insect diversity, the subtropics and tropics. As loss of seasonal patterns has been linked to a subsequent reduction in key ecosystem functions and services (Ross et al., 2021; Stevenson et al., 2015), increasing development could result in disruptions to nutrient cycling and food security (Bommarco et al., 2013). To better understand the generality of these patterns, future studies using high-frequency sampling should focus on monitoring a greater diversity of insect communities across different habitat strata (i.e., arboreal, subterranean; Gotelli et al., 2011) in a variety of biomes. Possibility widespread insect declines are receiving a great deal of attention, with good reason. We argue more attention on the impacts of land-cover change on insect community seasonality, and ecological seasonality more generally, should aid ongoing management and conservation efforts to help preserve the important ecosystem functions and services insects provide.

## Data Availability

Raw species’ count data before processing and R code used for analysis will be found on Data Dryad after article acceptance. Please contact <jamie.m.kass (at) gmail.com> with data requests.

## Acknowledgments

First and foremost, we would like to thank Takuma Yoshida and the OKEON Churamori Project Field Team for collecting, managing, and identifying the ant specimens: Linda Emiko Iha, Ayumi Inoguchi, Shinji Iriyama, Toshihiro Kinjo, Yoko Kudaka, Masafumi Kuniyshi, Izumi Maehira, Akino Miyagi, Seiichiro Nakagawa, Madoka Oguro, Shoko Suzuki, Yasutaka Tamaki, Takumi Uchima, and Kozue Uekama. We would also like to thank Samuel R. P. J. Ross and Nicholas R. Friedman for comments on earlier versions of the manuscript, and also Chisa Oshiro for administrative support. JMK was supported by the Japan Society for the Promotion of Science (JSPS) Postdoctoral Fellowships for Foreign Researchers Program and JSPS Grants-In-Aid for Scientific Research (KAKENHI) (20F20774). ID was supported by a JSPS Invitational Fellowship.

## Supporting Information

### A. Ant activity data details

As sites across the island were sampled in sequential groups rather than simultaneously for each biweekly period (differences are on the order of 7 days), to allow for appropriate site comparisons, we assigned each sample a time step (52 in total) based on the relative sampling order for that site. To this activity dataset, we filled in absence data when species that were observed at least once at a site were not observed for other time steps for that site in order to calculate temporal variability metrics. Values of NA (*n* = 8) were assigned to samples that experienced trap malfunctions or data loss via extreme weather events and were removed from analyses.

### B. Rarefaction permutations

Community samples collected over time are typically biased by sampling error, which results in fluctuations in activity due to site placement or other external factors, but this bias can be reduced via data rarefaction (Gaston & McArdle, 1994). As natural differences in total community count across sites were high (from *n* = 3,616 [Yona Forest] to *n* = 82,781 [Sueyoshi Forest]), we rarefied site- and group-level count while retaining original activity patterns to reduce the effects of sampling error, allowing us to make more even comparisons of temporal variability. For site-level activity, we randomly resampled the individual counts of species *i* at time *t* for each site to equal the minimum total site count (*n* = 3,616). For group-level activity, we first limited the species considered to those with total count greater than 100 for the two-year study period, then randomly resampled (with replacement) the individual counts of species *i* at time *t* for each group to equal that species’ minimum total count between groups. We performed 1000 iterations of each rarefaction and conducted all proceeding analyses on these iterations.

### C. Community and compositional variability calculations

We measured community temporal variability by calculating the coefficient of variation (CV) of summed ant count by site over the 2-year sampling period. We measured compositional temporal variability by calculating temporal beta diversity for each site community as the total variability of the species composition matrix (here, species × time step) after a Hellinger transformation (to ensure purely relative count data; Legendre & De Cáceres, 2013) using the function *beta.div* from the R package *adespatial* (Dray et al., 2022). As an additional product of this function, we derived species contributions to beta diversity per site, which is calculated as species variance divided by the total community variance (summing to 1). We then summed these contributions for each invasive status category (native, uncertain, alien) to determine the contribution of each category. We calculated both indices of temporal variability on each rarefied site activity dataset, then found the mean values across datasets. These mean rarefied site values were used in subsequent models.

### D. Formulas for temporal decomposition and associated variance

We used the TSLM() function in the R package *fable* (O’Hara-Wild et al., 2021) to fit time-series linear models with the following formula:

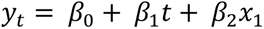

where *y*_t_ is the total count of ants at a site, with *t* equal to the range 1 to *T*, the number of time intervals; and where *x_1_* is the 1^st^ order Fourier term 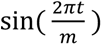, with *m* equal to the number of seasonal periods (in this case, *m* = 2 for 2 years).

We calculated the temporal components in the following ways:

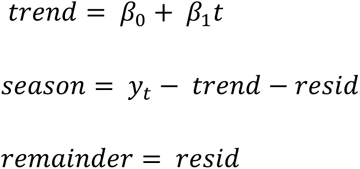

where *resid* is the model residuals.

In addition to calculating absolute variance of temporal components, we calculated relative variance with the following formula:

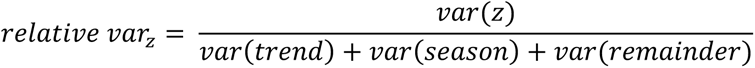

where *z* is one of the temporal components: trend, season, or remainder.

### E. Assessing effects of spatial autocorrelation on models

We first built models using ordinary least squares (OLS) regression to estimate the relationships between land cover and the variables of interest. As some OKEON sites are relatively close to others, we assessed the spatial autocorrelation present in the OLS model residuals by calculating Moran’s I and testing with random permutations using the function *moran.randtest*() from the R package *adespatial* (999 repetitions; Dray et al., 2022). But as this test was not significant for any of the models (*p* > 0.05), we fitted OLS models instead of more complex generalized least squares (GLS) models with spatial structure. We determined goodness-of-fit with the adjusted coefficient of determination (R-squared) and compared these between different count threshold choices.

### F. Effects of count threshold choice

As we avoided bias in temporal variability estimates by limiting all count values to an assigned threshold, we examined how robust our results are over a range of different thresholds. We report results in the main text for the relatively conservative threshold of 500 set at the station- level, but additionally ran the same analyses with no threshold, and with thresholds 100, 200, 1000, and 2000, applied at both the station- and site-levels. We observed some differences across the different thresholding schema, but no threshold fundamentally changes our results (which we report for threshold 500 set at the station-level). The adjusted R-squared values for the metrics calculated were sensitive in varying degrees to threshold (Fig. S7), with the exception of compositional temporal variability—this is because our beta diversity calculation includes a Hellinger transformation that mollifies the effects of outliers in the data. Further, thresholding at the station- and site-levels showed the same general trends. The highest threshold considered (2000) for the station-level was most often associated with the lowest R-squared values. The absence of a threshold had the most dramatic differences. This reduced the R-squared value for functional temporal variability to 0.11 from a thresholded minimum of 0.40, and absolute seasonality variance to 0.16 from 0.39. R-squared values for the other metrics were reduced to 0 when thresholding was not used. These reductions in correlations with PC1 occurred because, without thresholding, sites with extreme count values for one or several species spike in functional variability (calculated with CV) and lose seasonality, and the existence of such spikes has little to do with land cover.

**Table S1.** Pearson correlations between proportions of land-cover classes within 1 km buffers of sampling sites. “MISC” stands for miscellaneous and includes the classes for unclassified pixels and those classified as bare rock or soil. As pairwise correlations were high, we conducted a principal component analysis (PCA) to derive the main orthogonal land-cover gradients.

|  | AGRICULTURE | FOREST | WATER | GRASS | SAND | URBAN | MISC |
| --- | --- | --- | --- | --- | --- | --- | --- |
| AGRICULTURE | 1 |  |  |  |  |  |  |
| FOREST | -0.89 | 1 |  |  |  |  |  |
| WATER | -0.05 | 0.02 | 1 |  |  |  |  |
| GRASS | 0.53 | -0.44 | 0.02 | 1 |  |  |  |
| SAND | 0.54 | -0.48 | -0.04 | 0.23 | 1 |  |  |
| URBAN | 0.56 | -0.84 | -0.08 | 0.01 | 0.20 | 1 |  |
| MISC | 0.51 | -0.73 | 0.13 | 0.18 | 0.53 | 0.74 | 1 |

**Table S2.**
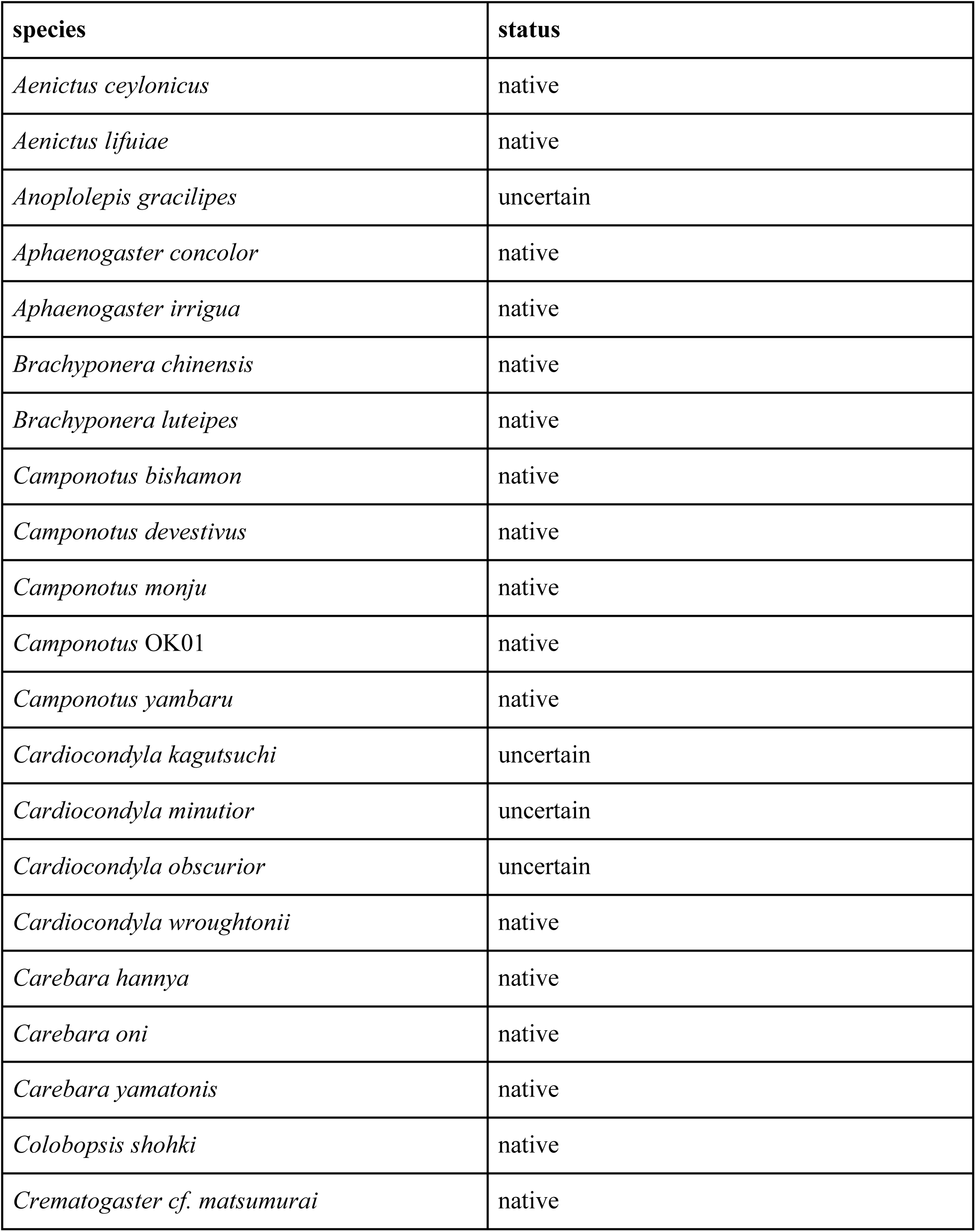
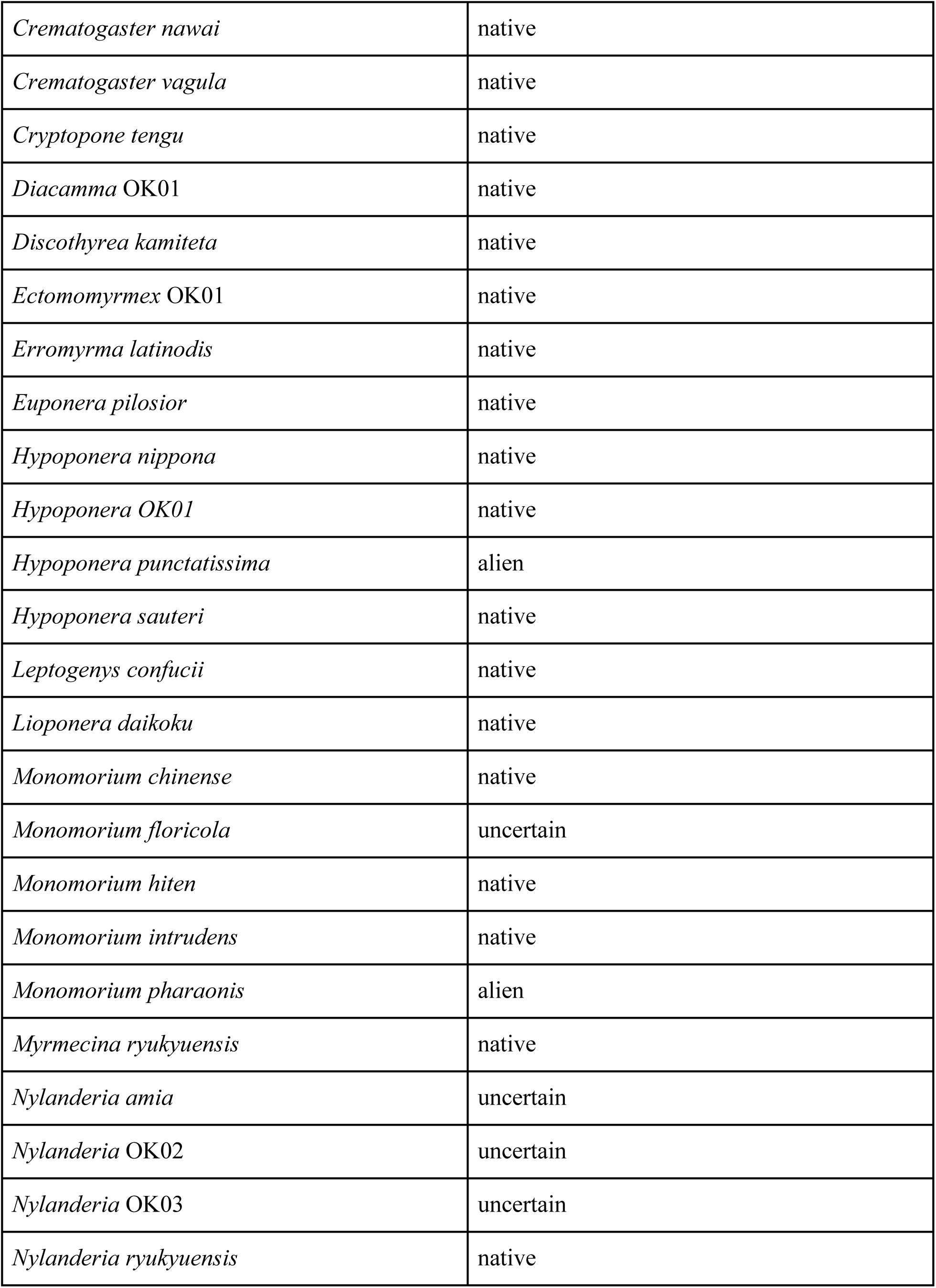
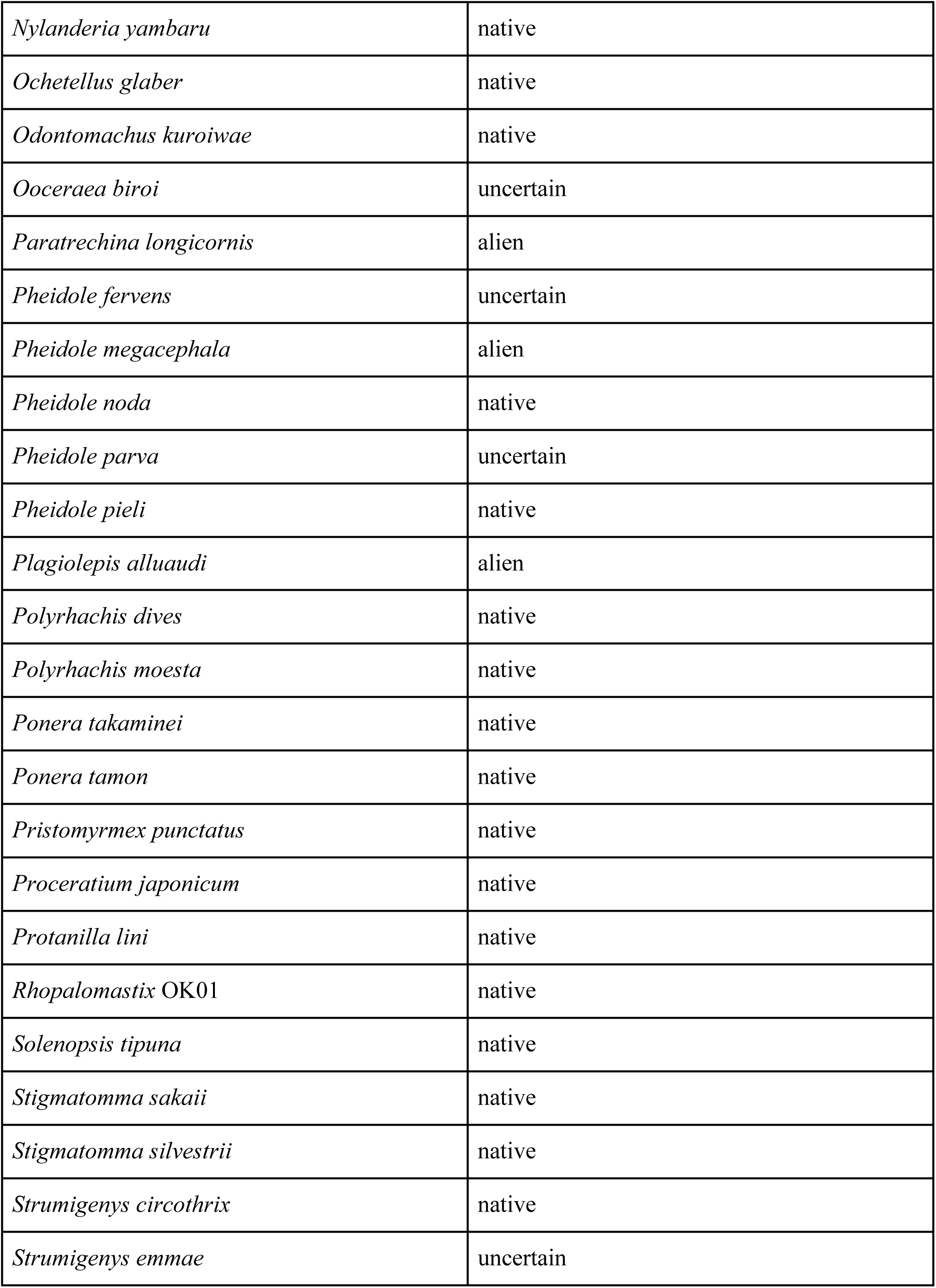
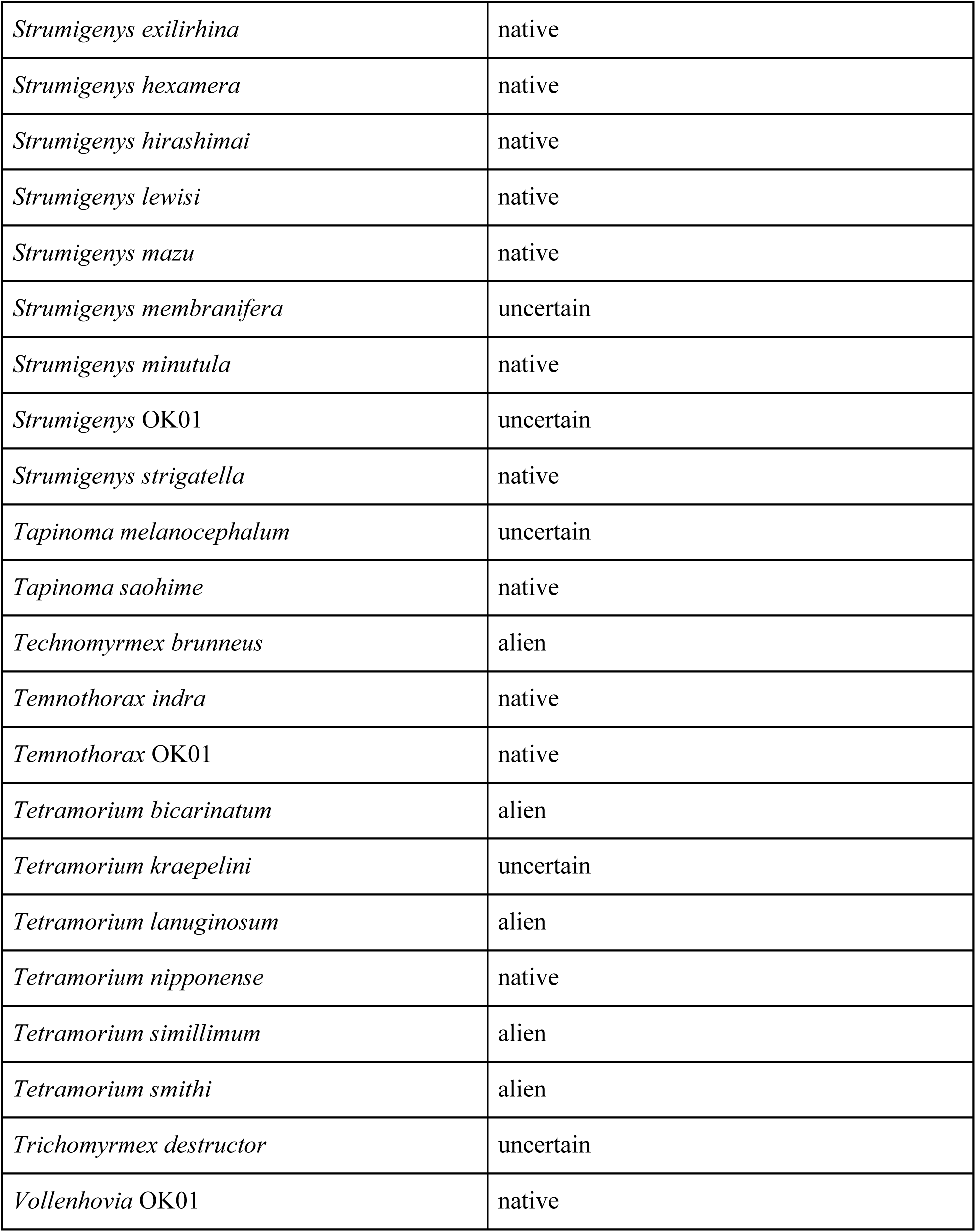
Invasive status for all observed ant species based on existing literature and expert opinion.

**Table S3.** Results from linear regression models built with two land-cover PCA axes: PC1, explaining the forested-developed gradient, and PC2, explaining the rural-urban gradient. All model residuals were tested for spatial autocorrelation by calculating Moran’s I and testing with random permutations (*moran.randtest*() from the R package *adespatial*; 999 repetitions), yet none had significant results.

|  |  | PC1 |  |  | PC2 |  |  | Adj.<br>R<sup>2</sup> | Moran<br>*p* |
| --- | --- | --- | --- | --- | --- | --- | --- | --- | --- |
| Response variable | Intercept | Estimate | Std. Error | *p* | Estimate | Std. Error | *p* |  |  |
| Total activity (natural log) | 10.082 | -1.575 | NA | 0.206 | NA | 0 | NA | 0.714 | 0.279 |
| Total richness (rarefied) | 22.753 | 3.484 | NA | 1.558 | NA | 0.036 | NA | 0.148 | 0.753 |
| Native richness (rarefied) | 13.67 | 8.363 | NA | 1.368 | NA | 0 | NA | 0.612 | 0.803 |
| Uncertain richness (rarefied) | 5.864 | -3.573 | NA | 1.022 | NA | 0.002 | NA | 0.328 | 0.157 |
| Alien richness (rarefied) | 3.219 | -1.306 | NA | 0.507 | NA | 0.017 | NA | 0.197 | 0.593 |
| Total richness (observed) | 30.708 | NA | NA | NA | NA | NA | NA | 0 | 0.91 |
| Native richness (observed) | 18.708 | 7.483 | -7.594 | 1.686 | 4.611 | 0 | 0.114 | 0.47 | 0.783 |
| Uncertain richness (observed) | 7.875 | -5.188 | NA | 1.516 | NA | 0.002 | NA | 0.318 | 0.304 |
| Alien richness (observed) | 4.125 | -2.052 | NA | 0.643 | NA | 0.004 | NA | 0.285 | 0.269 |
| Total richness (estimated) | 38.923 | NA | NA | NA | NA | NA | NA | 0 | 0.887 |
| Native richness (estimated) | 23.781 | 13.457 | NA | 3.646 | NA | 0.001 | NA | 0.354 | 0.748 |
| Uncertain richness (estimated) | 8.427 | -5.717 | NA | 1.636 | NA | 0.002 | NA | 0.328 | 0.475 |
| Alien richness (estimated) | 4.292 | -2.092 | NA | 0.802 | NA | 0.016 | NA | 0.202 | 0.267 |
| Functional temporal variability | 0.67 | 0.405 | NA | 0.059 | NA | 0 | NA | 0.667 | 0.267 |
| Compositional temporal variability | 0.23 | 0.14 | NA | 0.024 | NA | 0 | NA | 0.601 | 0.721 |
| Native SCBD | 0.53 | 0.237 | NA | 0.059 | NA | 0.001 | NA | 0.398 | 0.832 |
| Uncertain SCBD | 0.229 | -0.16 | NA | 0.05 | NA | 0.004 | NA | 0.284 | 0.498 |
| Alien SCBD | 0.257 | -0.087 | NA | 0.05 | NA | 0.093 | NA | 0.083 | 0.46 |
| Seasonal variability (absolute) | 1208.507 | 1851.371 | NA | 328.794 | NA | 0 | NA | 0.572 | 0.925 |
| Seasonal variability (relative) | 0.478 | 0.201 | -0.375 | 0.081 | 0.222 | 0.022 | 0.106 | 0.233 | 0.389 |
| Trend variability (absolute) | 194.827 | 311.661 | NA | 90.442 | NA | 0.002 | NA | 0.321 | 0.546 |
| Trend variability (relative) | 0.079 | NA | NA | NA | NA | NA | NA | 0 | 0.692 |
| Stochastic variability (absolute) | 946.779 | 660.402 | NA | 297.651 | NA | 0.037 | NA | 0.146 | 0.604 |
| Stochastic variability (relative) | 0.443 | -0.223 | 0.347 | 0.066 | 0.18 | 0.003 | 0.068 | 0.364 | 0.412 |

**Figure S1.**
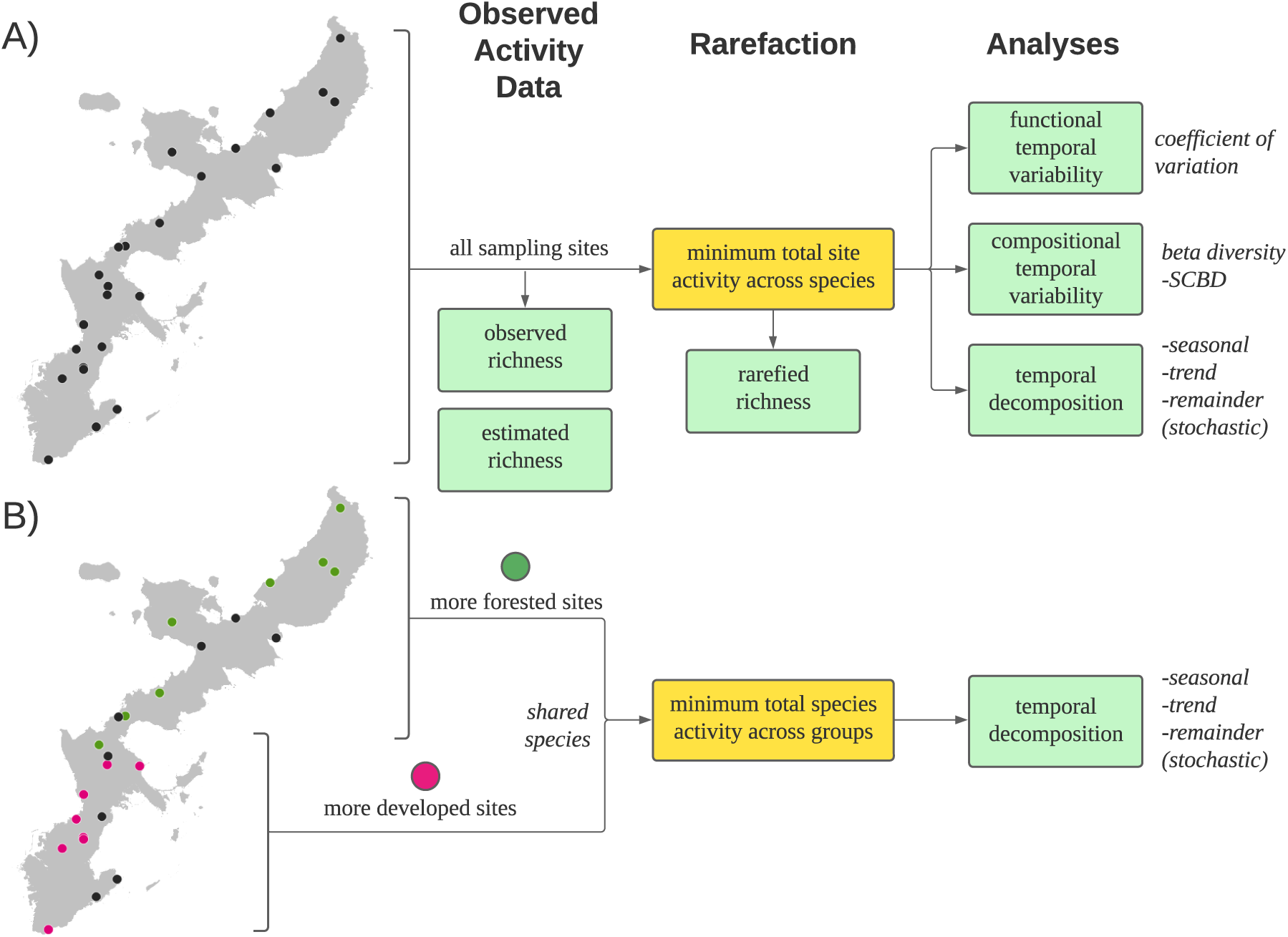
Conceptual workflow schematic describing the analysis steps. A) The main analysis uses ant activity data from all sites. Richness is calculated for observed data and estimated via Hill numbers, then the dataset is rarefied by randomly resampling individual workers per site to match the minimum total site activity; rarefied richness is calculated from this rarefied dataset. Next, the following analyses are conducted: functional temporal variability, compositional temporal variability (including species contribution to beta diversity [SCBD]), and temporal decompositions to derive seasonal, trend, and remainder (stochastic) components. B) The group-level analysis first splits sites into two groups (green and pink points) based on land-cover PC1, explaining the forested-developed gradient. Both groups share the same species, and those with mid-level PC1 values are excluded (black points). These grouped data are then rarefied by randomly resampling individual workers per species within each group to match the minimum total species activity between groups. The same temporal decomposition is then conducted on these grouped data.

**Figure S2.**
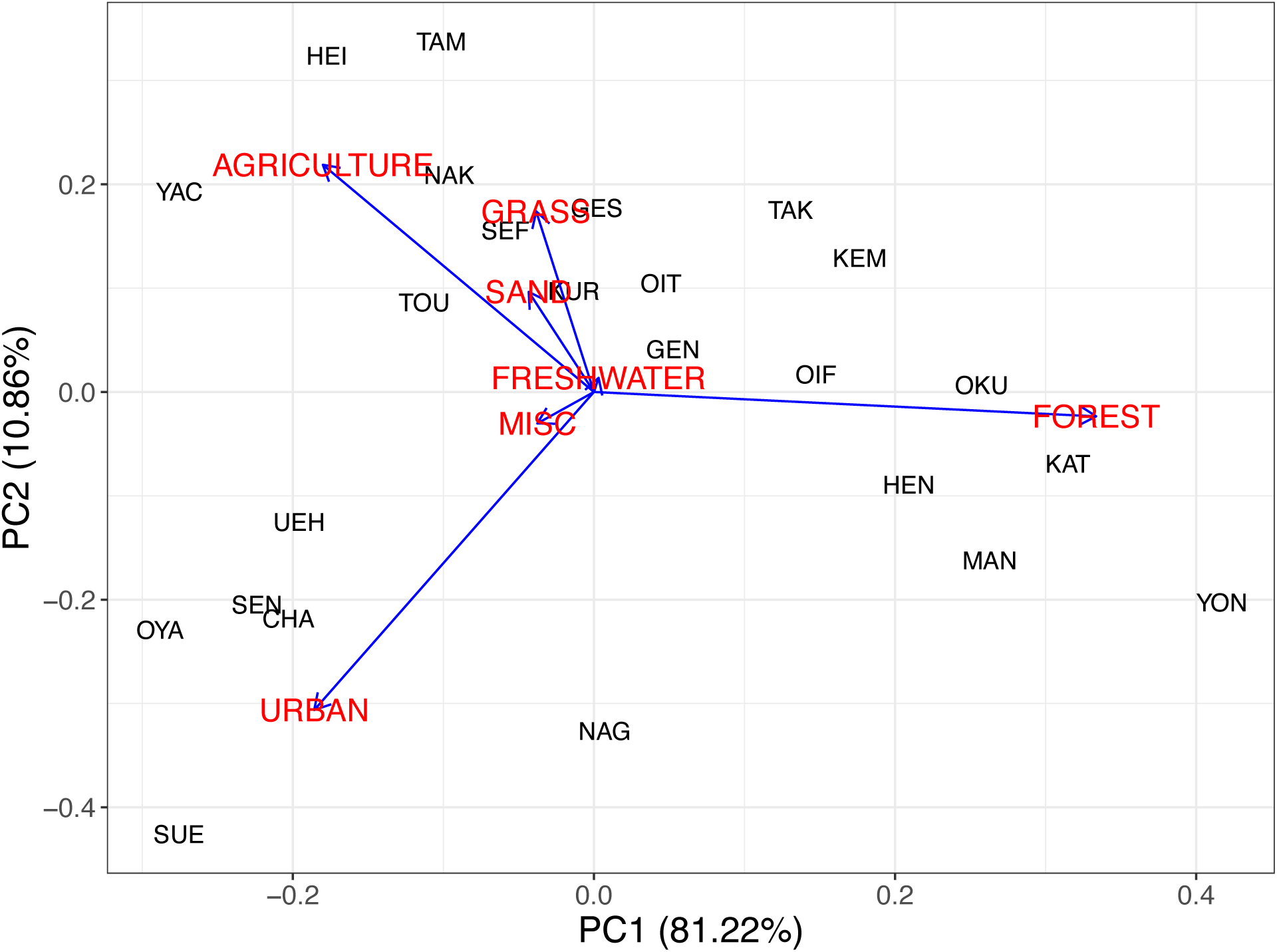
Biplot of the principal component analysis for the land-cover proportion dataset. PC1 explains the main anthropogenic stress gradient (low: more forested, high: more developed), while PC2 explains the rural (high; agriculture and/or grass) to urban (low) gradient.

**Figure S3.**
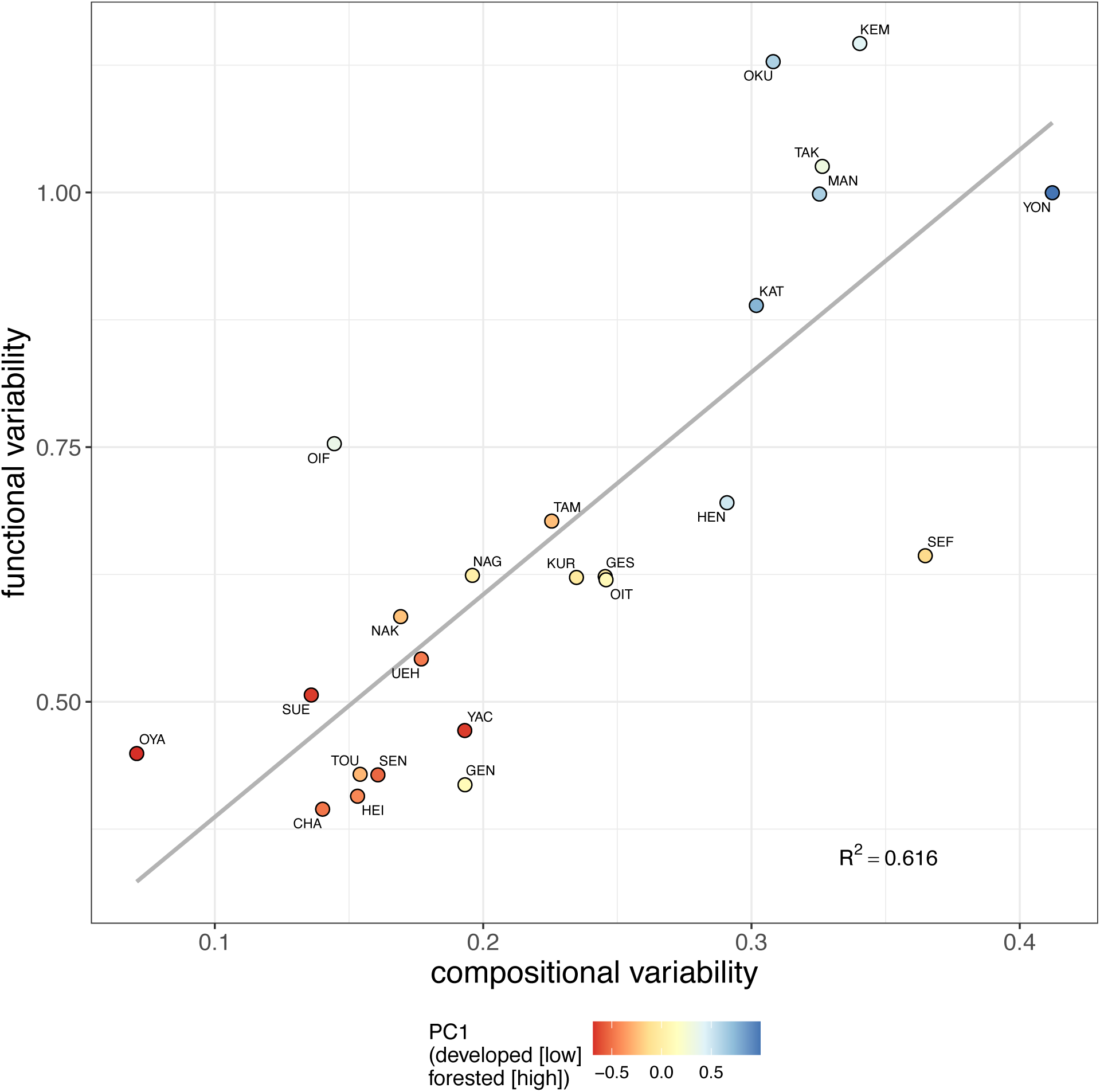
Plot of compositional variability by functional variability, showing the sampling sites with colors corresponding to the forested-developed gradient (PC1), where higher values (blue) are more forested and lower values (red) have more human development. Site abbreviations correspond to those used in Figure 1. The gray line shows the relationship between two variabilities modeled with linear regression.

**Figure S4.**
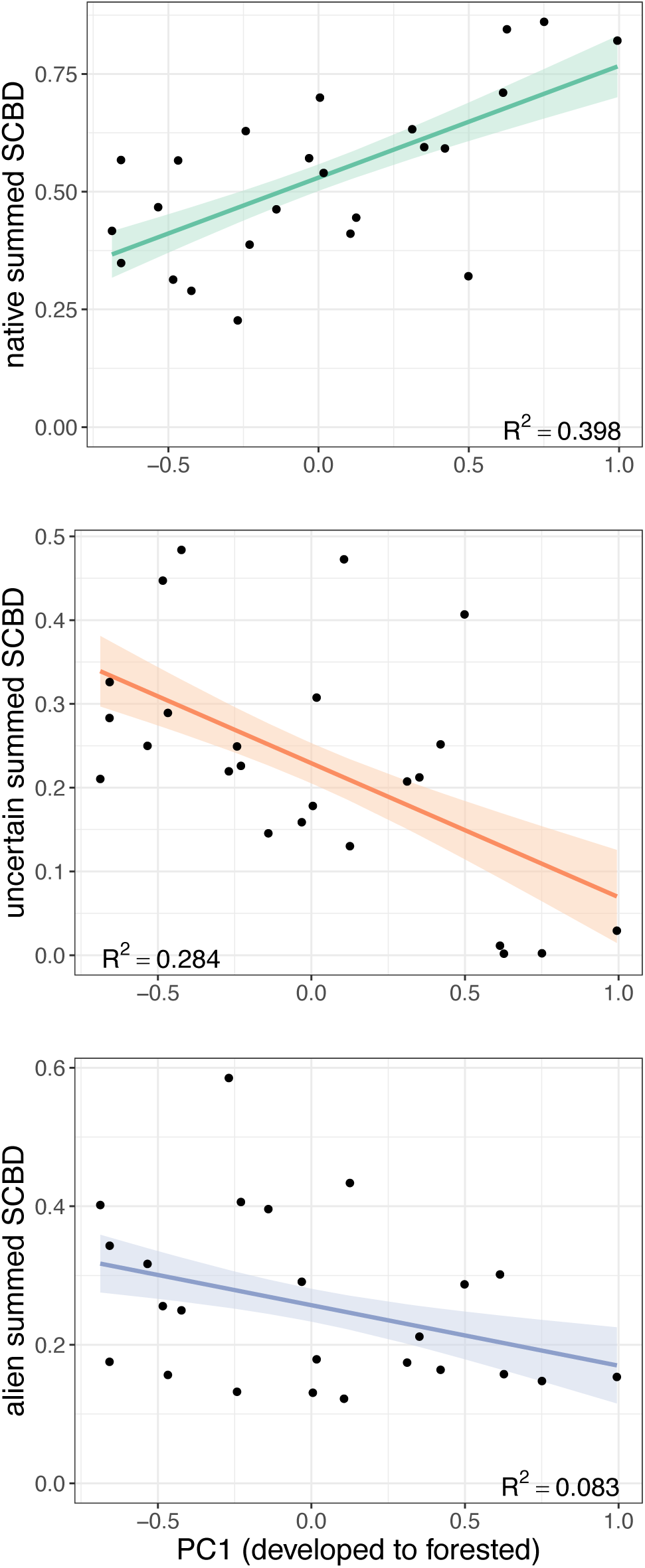
Relationships between PC1 and summed species contributions to beta diversity (SCBD) for native, uncertain, and alien species for the rarefied activity data. PC1 explains the anthropogenic stress gradient (low: more developed, high: more forested).

**Figure S5.**
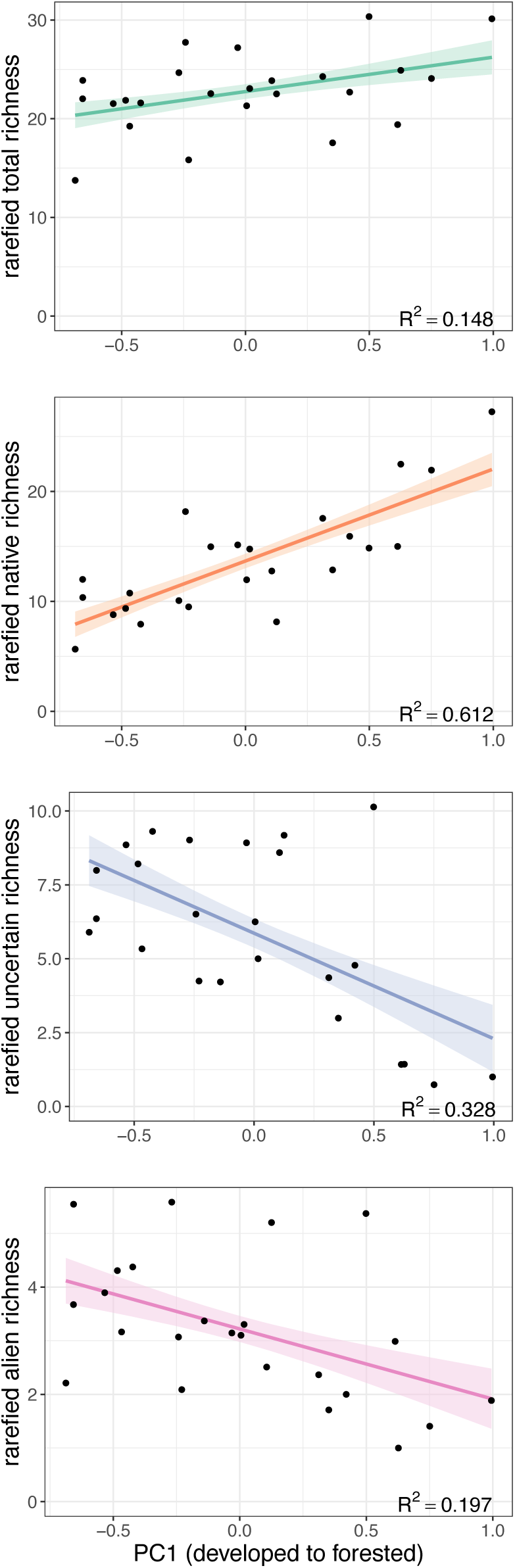
Relationships between PC1 and total richness, as well as richness of native, uncertain, and alien species for the rarefied activity data. PC1 explains the anthropogenic stress gradient (low: more developed, high: more forested).

**Figure S6.**
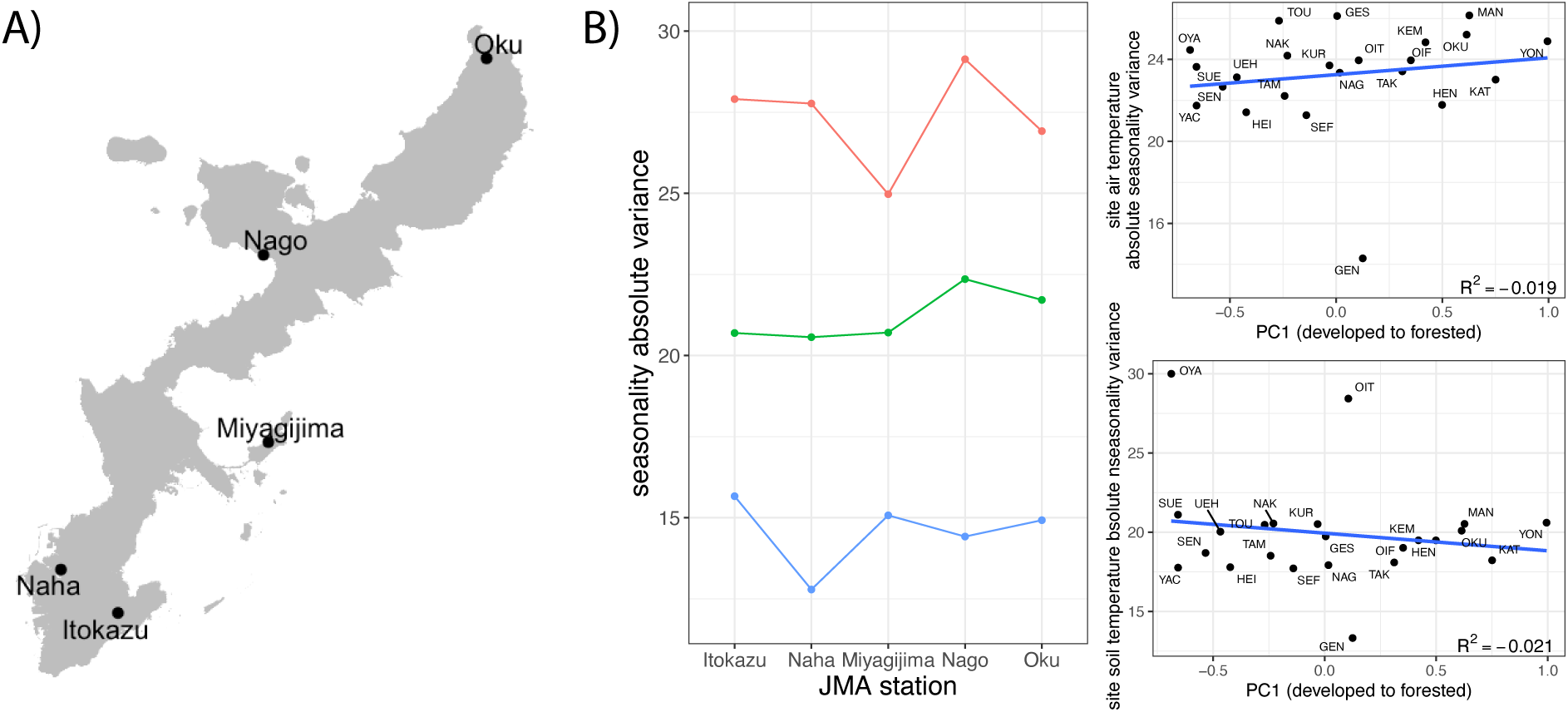
Absolute seasonality variance of temperature at the a) regional and b) site-level scales over the two-year study period (2016 – 2018). Regional temperature measurements were taken from Japanese Meteorological Association weather stations, and station names correspond to those on the map. Site-level temperature measurements were taken for air and soil at one station per sample site. Oyama Park (OYA) and OIST Open (OIT) were the only sites with sensors not located below forest canopy, and thus have higher variance for soil temperature seasonality.

**Figure S7.**
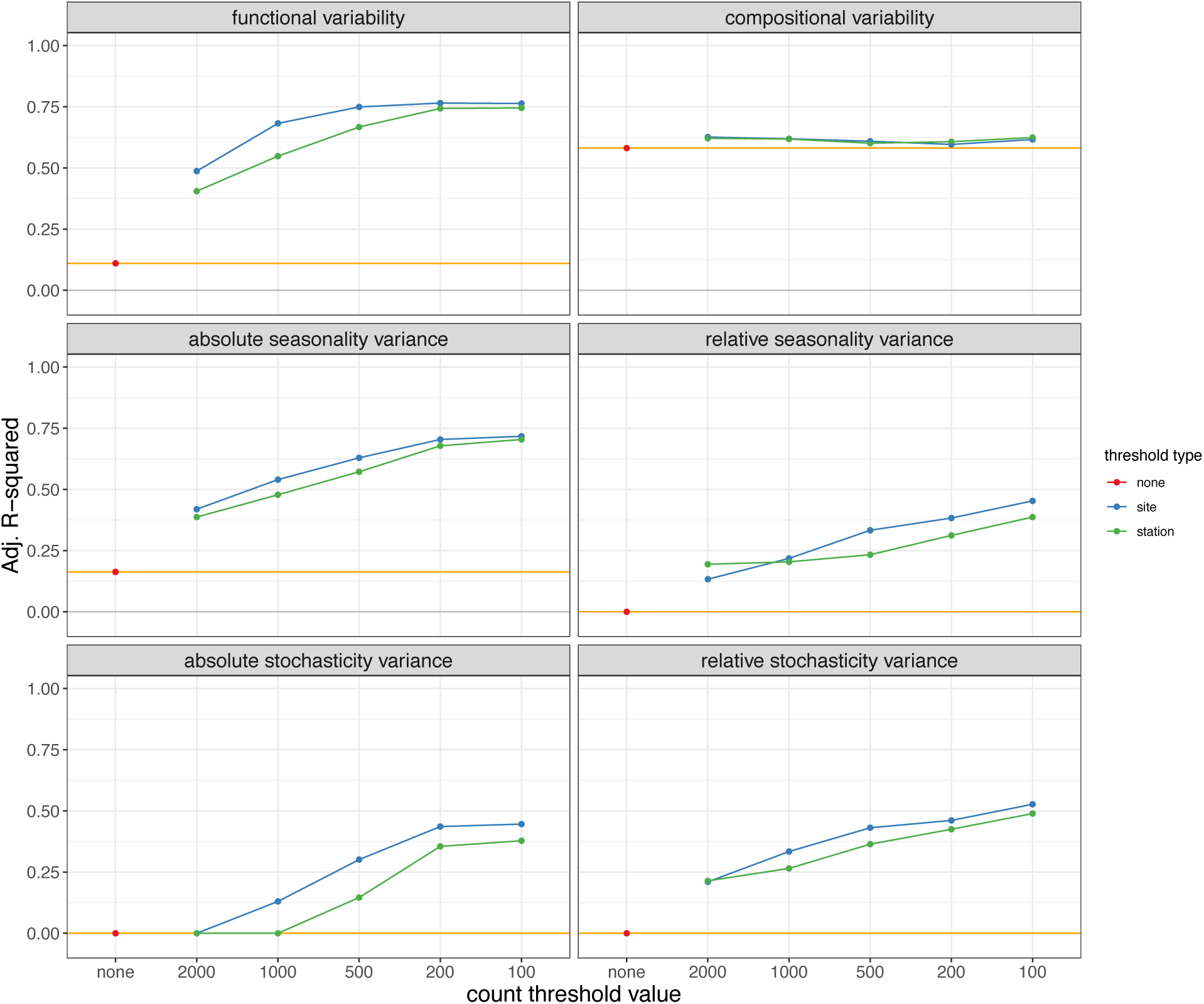
The adjusted R-squared value for the metrics measured in this study at different count thresholds and spatial levels for thresholding (station, site). The threshold value “none” is used as a reference and is symbolized by the red point and orange line. For reference, the analysis described in this paper is a threshold value of 500 at the station level.

## Notes

### Competing Interest Statement

The authors have declared no competing interest.

### Summary of Updates

We made some significant textual changes to the manuscript based on reviews that added background information and more nuanced discussion about the time-scale of the study and the sampling methods used. No results have changed.

